# Actin-binding compounds, discovered by FRET-based high-throughput screening, differentially affect skeletal and cardiac muscle

**DOI:** 10.1101/2020.05.19.104257

**Authors:** Piyali Guhathakurta, Lien A. Phung, Ewa Prochniewicz, Sarah Lichtenberger, Anna Wilson, David D. Thomas

**Author notes:** Authors equally contributed to manuscript (Piyali Guhathakurta, Lien A. Phung). Corresponding author: David. D. Thomas.

## Abstract

We have used spectroscopic and functional assays to evaluate the effects of a group of actin-binding compounds on striated muscle protein structure and function. Actin is present in every human cell, and its interaction with multiple myosin isoforms and multiple actin-binding proteins is essential for cellular viability. A previous high-throughput time-resolved fluorescence resonance energy transfer (TR-FRET) assay from our group identified a class of compounds that bind to actin and affect actomyosin structure and function. In the current study, we tested their effects on the two isoforms of striated muscle α-actins, skeletal and cardiac. We found that a majority of these compounds affected the transition of monomeric G-actin to filamentous F-actin, and that these effects were different for the two actin isoforms, suggesting a different mode of action. To determine the effects of these compounds on sarcomeric function, we further tested their activity on skeletal and cardiac myofibrils. We found that several compounds affected ATPase activity of skeletal and cardiac myofibrils differently, suggesting different mechanisms of action of these compounds for the two muscle types. We conclude that these structural and biochemical assays can be used to identify actin-binding compounds that differentially affect skeletal and cardiac muscles. The results of this study set the stage for screening of large chemical libraries for discovery of novel compounds that act therapeutically and specifically on cardiac or skeletal muscle.

## Introduction

Muscle contraction results from interaction between myosin and actin, in which the transition from relaxation to contraction depends on both (a) calcium-mediated changes in the actin-bound regulatory proteins in the troponin complex (TnI, TnC, TnT) and tropomyosin, and (b) additional cooperative activation due to the strong binding of myosin to actin (1). The sarcomere is in a dynamic equilibrium between relaxed and contracted states, with calcium concentrations shifting the balance. Under resting conditions with low calcium, the troponin complex (TnT, TnI, TnC) maintains actin–tropomyosin in a ‘blocked’ state that prevents actin–myosin interaction (2). When the intracellular calcium concentration increases, a conformational change in the troponin complex allows tropomyosin to shift into the ‘open’ position that promotes strong actin–myosin binding (2), thereby triggering muscle contraction mediated by the chemical energy generated from myosin ATP hydrolysis (Fig. 1). Genetic mutation or a post-translational modification (e.g., oxidation, glycation) of a sarcomeric protein can alter its structure and function and have a detrimental effect on the process of contractility (3,4). The binding of a small molecule to that altered protein can restore normal contractility (5). Such therapeutic molecules or drugs can work indirectly by intervening in signaling pathways, or directly through interaction with contractile proteins (6). Discovery of such modulators presents a novel approach to treating any condition in which striated muscle function is compromised, including heart failure, cardiomyopathies, skeletal myopathies and a wide range of neuromuscular conditions.

**Figure. 1.**
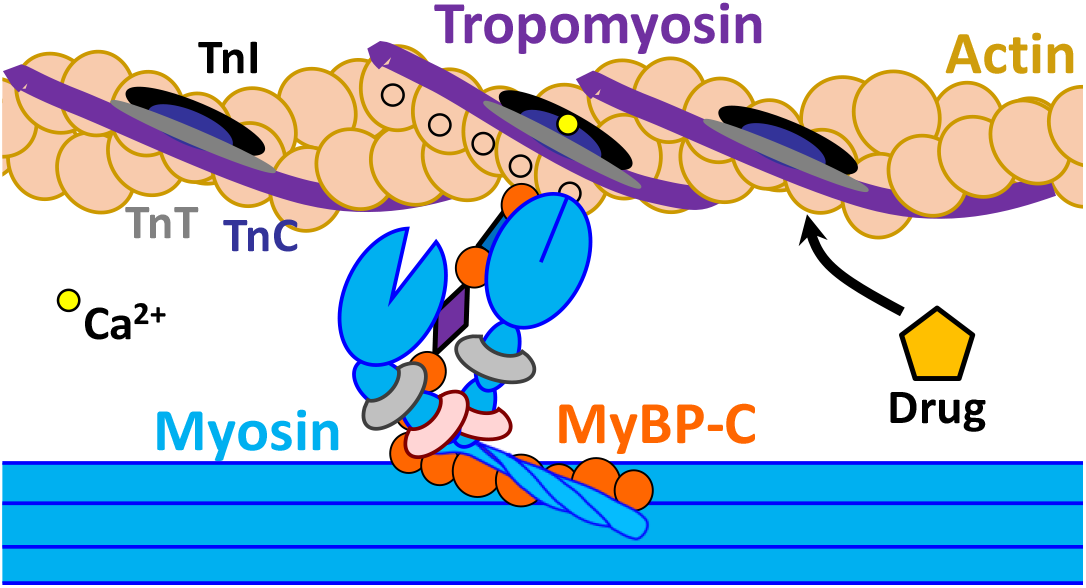
Schematic representation of thin and thick filaments substructures, key participants in Ca^2+^-dependent muscle contraction and relaxation. Actin-binding compounds can play a crucial role in regulation of these processes.

Sarcomere modulators identified through small-molecule screening have been shown to act at three different levels: (a) signaling pathways that affect protein phosphorylation, (b) direct interaction with myosin and (c) interaction with the troponin complex (6). Small-molecule effectors designed to target and modulate striated and smooth muscle myosin isoforms for the treatment of diseases are showing some promising results in preclinical and clinical trials. A cardiac myosin effector, Omecamtiv Mecarbil (OM), identified via a low-throughput functional assay, has so far only marginally improved the condition of heart failure patients compared to placebo controls (7). Another cardiac myosin modulator, Mavacamten (Myk461), was found to decrease cardiac contractility; and in the future may be applied to treat diseases in which hypercontractility is an issue (8). Several troponin-specific modulators have been identified. Tirasemtiv and CK-2066260, the most advanced of these agents, binds specifically to fast skeletal troponin C (9,10) and has no observable affinity for cardiac troponin C. In drug development for muscle disorders, it is critical to determine a compound’s specificity for muscle type in a high-throughput manner, so that we can maximize its advantages and minimize its disadvantages in clinical application.

None of the previous studies attempted to target striated muscle actin, the main binding companion of striated myosin. Actin is a highly conserved protein and interacts with multiple proteins in both muscle and non-muscle cells. Actin is flexible and dynamic, and its different structural states can be cleverly sensed by its binding partners. Higher eukaryotes have six different actins, expressed from separate genes (12), with most variability between the proteins occurring at their N-terminal region. In this work, we focus on skeletal and cardiac α-actin. These two isoforms are 99% homologous, with only 4 amino acid substitutions from skeletal to cardiac: Glu2Asp, Asp3Glu, Met299Leu, and Thr358Ser (13); nevertheless, the two isoforms show significant differences in their functional properties (13,14). The rates of actin polymerization and activation of myosin ATPase are essentially the same among these actin isoforms (15), but myosin cross-bridge formation and myofiber force production are significantly different (16). In their respective muscles, skeletal and cardiac actin filaments interact with myosin and regulatory proteins such as troponin, tropomyosin, and myosin binding protein C (MyBP-C), at different stages of contraction and relaxation. These interactions are finely tuned, and we expect that a subtle change by actin-binding modulators can play an important role in therapeutic treatment for muscle disorders (Fig. 1). Recent studies from our group utilized a high-precision time-resolved fluorescence resonance energy transfer (TR-FRET) approach for high-throughput screening (HTS) of a small-molecule library and detected several compounds that bind to actin and affect actomyosin structure and function (11) (Table 1). In the current study, we have established reliable biochemical and structural assays to test whether these compounds are specific for skeletal or cardiac muscle protein isoforms. In particular, we focus on the effects on polymerization of both actin isoforms, as well as interaction with myosin in the presence of regulatory proteins, as detected by myofibrillar ATPase activity. We found that these compounds have subtle but differential effects on different isoforms of actin, which impact function of the corresponding muscle type. Thus, our results set the stage to screen large chemical libraries for discovery of novel actin-binding drugs that act specifically on cardiac or skeletal muscle.

**Table 1.** Actin-binding compounds detected from high-throughput FRET assay^(11)^

| Compound | Clinical Use |
| --- | --- |
| 1. Honokiol | anticancer |
| 2. Fluphenazine | antipsychotic |
| 3. Phenothiazine | antipsychotic |
| 4. Tegaserod | irritable bowel syndrome |
| 5. Carvedilol | $\beta$ blocker, treats high BP |
| 6. Novantrone | multiple sclerosis |
| 7. Thioridazine | antipsychotic |
| 8. Flutamide | nonsteroidal antiandrogen anticancer |
| 9. Mefloquine | anti-malaria |
| 10. Dantrolene | postsynaptic muscle relaxant |

## Results

### Effects of compounds on actin polymerization and depolymerization

The time course of polymerization of rabbit skeletal and bovine cardiac G-actin was quantified by the increase in fluorescence intensity of pyrene-labeled actin with excitation at 350 nm and emission at 407 nm (Fig. 2). When pyrene-G-actin polymerizes to form F-actin, the pyrene fluorescence intensity increases 7-10 fold, in proportion to the fraction of G-actin incorporated into F-actin (17). Control experiments showed that the compounds in Table 1 did not have any effect on the fluorescence of unlabeled G-actin. The polymerization behavior of skeletal and cardiac actin was first monitored in the presence of either 50 mM KCl or 2 mM MgCl_2_. The polymerization rate is faster in the presence of MgCl_2_ for both isoforms (Fig. 2), as reported earlier (17),. The effects of the compounds (listed in Table 1) on polymerization of both actin isoforms were then determined in the presence of 50 μM compound (Fig. 2, Fig. 3). Prior to the experiment, skeletal and cardiac G-actin containing 5% pyrene G-actin were incubated with compounds for 10 minutes at 23 °C. After incubation, 2 mM MgCl_2_ was added, and polymerization was monitored for at least 15 minutes, until the saturation of fluorescence intensity was achieved.

**Figure. 2.**
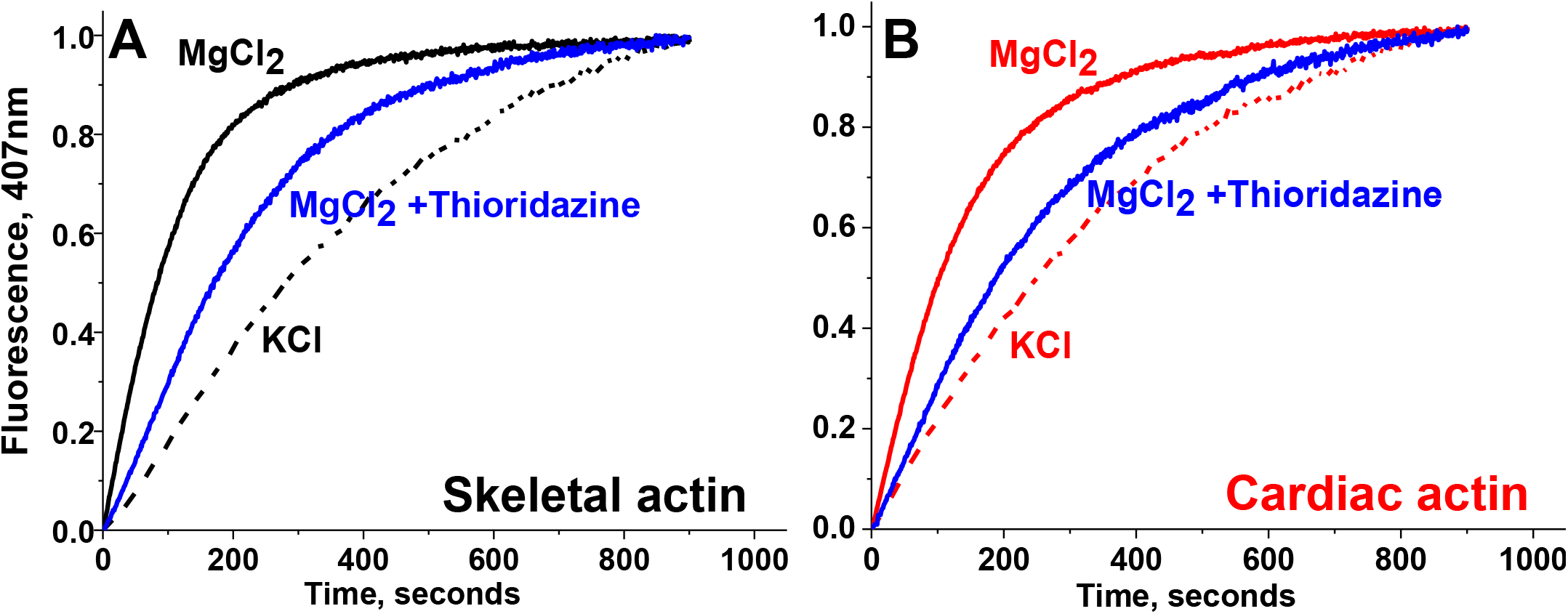
Salt and compound sensitivity of actin polymerization. (A) Rabbit skeletal actin. (B) Bovine cardiac actin. In both cases polymerization in faster in the presence of MgCl_2_ (2 mM) than in the presence of KCl (50 mM). Addition of 50 μM thioridazine slows polymerization of both actin isoforms.

**Figure. 3.**
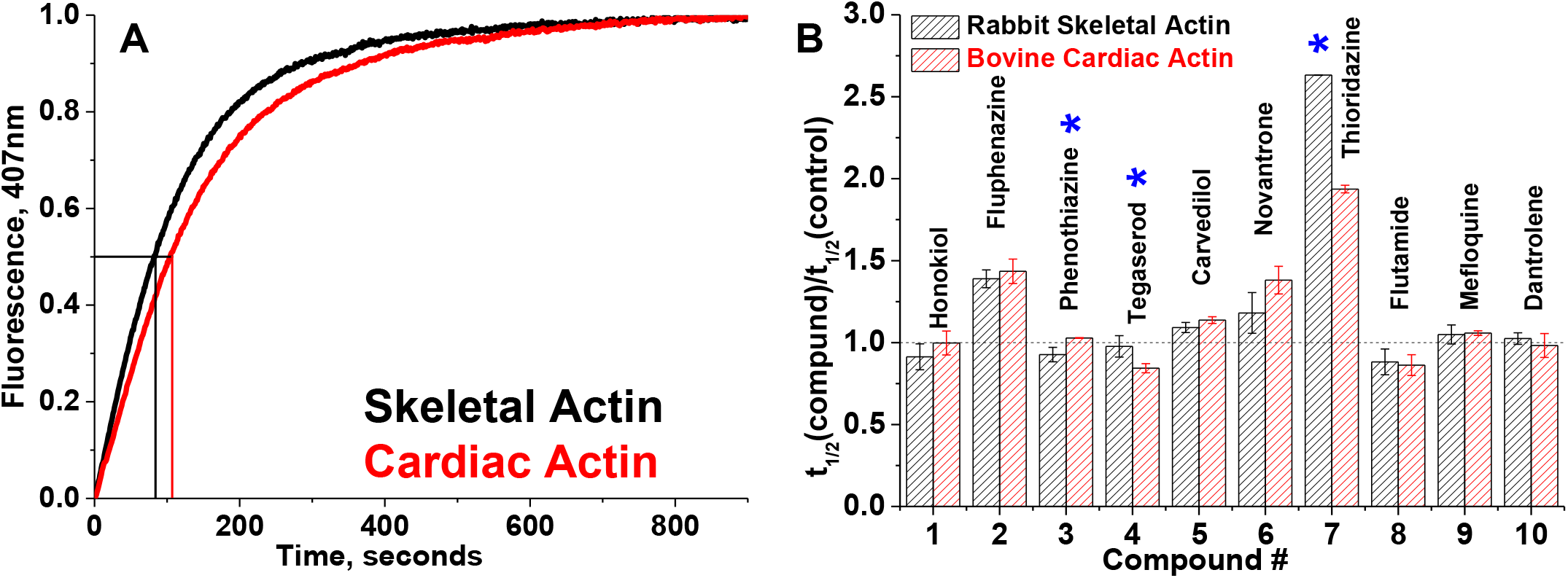
Actin polymerization. (A) Time courses of polymerization of skeletal and cardiac actin in the absence of compounds. The increase in fluorescence of skeletal and cardiac actin, containing 5% pyrene-actin, was monitored at 407 nm. Polymerization half-times (vertical lines) denote the times required for actin to reach half maximal fluorescence (horizontal line). For each actin species, at least three independent preparations were used. (B) Relative change in apparent half-times (t_1/2_) of polymerization of actin isoforms in the presence of 50 μM compound. Errors are reported as standard deviation, n = 3. Significance (*) of difference in effects between skeletal and cardiac was established with Student’s t-test, P<0.05.

In the absence of compounds, both skeletal and cardiac actin show similar polymerization behavior (Fig. 3A). The effect of compound on actin isoform was determined from the ratio of polymerization half time (t_1/2_) in the presence and absence of compound (Fig. 3B). Fluphenazine, carvedilol, novantrone and thioridazine reduce the rate of polymerization (higher t_1/2_) of both actin isoforms. However, with thioridazine, the decrease is more pronounced for skeletal compared to cardiac. Honokiol and phenothiazine slightly increase the rate of polymerization of skeletal actin. Tegaserod and flutamide increase the rate of polymerization of both actin isoforms, but the effect is more pronounced on cardiac actin. Dantrolene and mefloquine did not have a significant effect on the rate of polymerization of either skeletal or cardiac actin. Thus most of these compounds affect actin polymerization in an isoform-specific manner. These alterations are subtle but significant and could be related to changes in actin’s functional interaction with myosin and regulatory proteins.

To evaluate the effects of drugs on actin depolymerization, 20 μM actin containing 5% pyrene actin was first polymerized with 2 mM MgCl_2_ for at least 2 hours at 23°C. This F-actin was diluted 1:20 into G-buffer and the decrease in fluorescence intensity was monitored at the wavelength settings indicated above. Figure 4 shows an example of depolymerization of cardiac actin in the presence and absence of a representative drug, thioridazine. The data indicates no change in depolymerization rate. None of these compounds had a significant effect on the rate of depolymerization of either actin isoform. These data suggest that both isoforms of actin are able to maintain filament integrity in the presence of compounds. Any observed functional alterations in muscle with the use of these compounds would not be due to the depolymerization of actin. This is encouraging, as depolymerization of muscle actin is not beneficial for muscle function. The significant effects on polymerization (Fig. 3) indicate perturbation of actin structure.

**Figure. 4.**
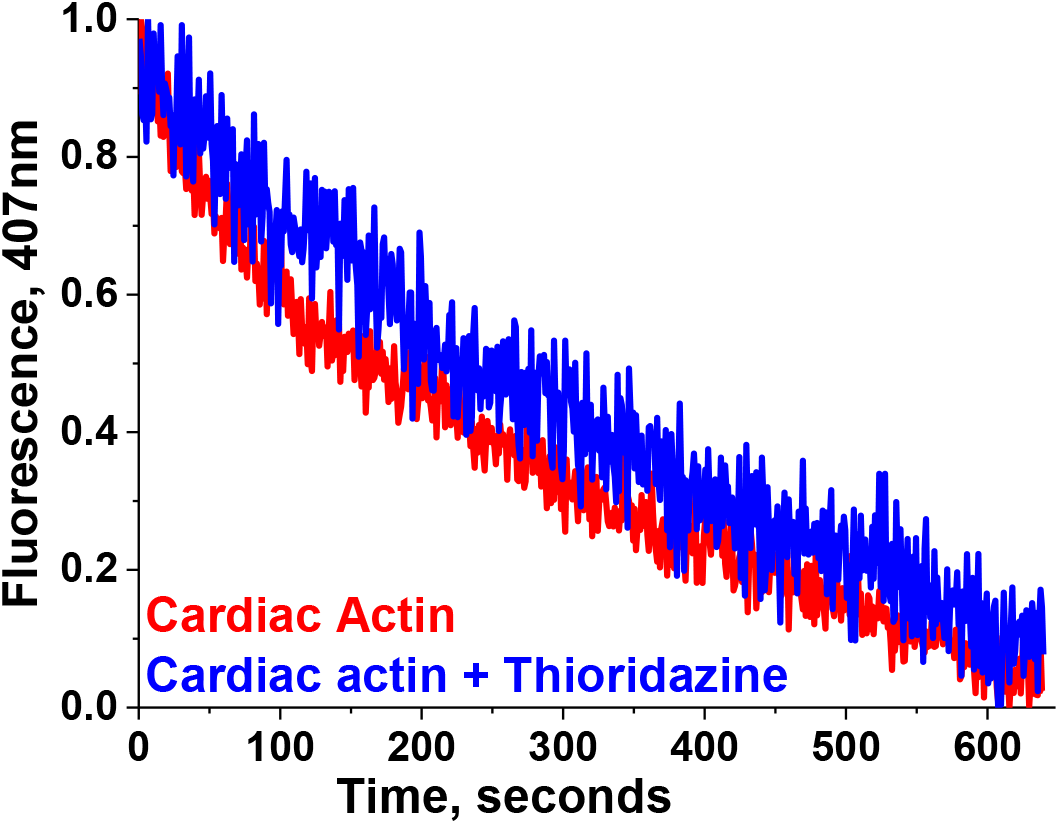
Effect of drug on depolymerization of F-actin. Cardiac F-actin depolymerization with and without 50μM Thioridazine

### Effects of compounds on skeletal and cardiac myofibril ATPase

To further determine the effects of compounds’ actin isoform specificity on muscle function, we examined the Ca^2+^-dependent ATPase activity of skeletal and cardiac myofibrils in a high-throughput manner, varying the free calcium concentration from pCa 8 (relaxation) to pCa 4 (full activation). Fig. 5A confirms that skeletal myofibrils have a 10-fold activation in response to calcium (10,18,19), whereas activation of cardiac myofibrils is 4-fold (20). At pCa 4, skeletal and cardiac myofibril ATPase show a V_max_ of 3.74 ± 0.22 s^−1^ and 0.31 ± 0.02 s^−1^, respectively. At pCa 8, their corresponding basal ATPase rates (V_0_) are 0.28 ± 0.01 s^−1^ and 0.08 ± 0.003 s^−1^. The pK_Ca_ values of skeletal and cardiac myofibrils are 6.06 ± 0.01 and 5.76 ± 0.02. These values are consistent with previous reports (10,11,18,20–22). The myofibril ATPase activities of both muscle systems were not affected either by the presence of 1% DMSO (data not shown) or by 500 nM thapsigargin, a SERCA (sarcoendoplasmic reticulum calcium transport ATPase pump) modulator (data not shown). Lack of any significant changes in the maximal ATPase activity or pK_Ca_ in either of the myofibril types indicates that the measured Ca^2+^-dependent activities are determined by myofilament based calcium regulation, and are not affected by contaminating residual SERCA in the preparation.

**Figure. 5.**
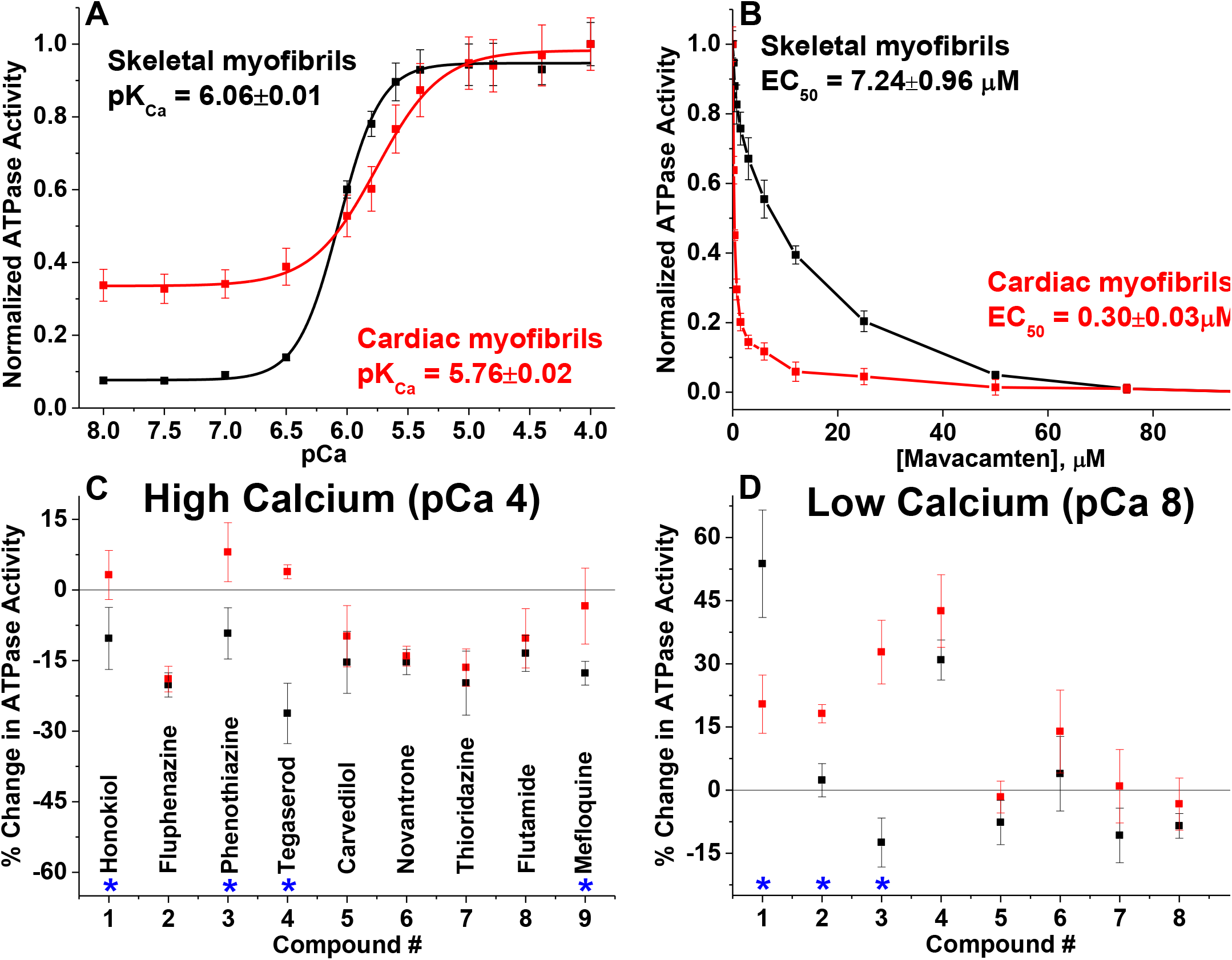
Effects of compounds on skeletal and cardiac myofibrillar ATPase activity at high and low calcium concentrations. (A) Ca^2+^-dependent ATPase activation of rabbit skeletal myofibrils (black, n = 8 independent preparations); and bovine cardiac myofibrils (red, n = 10 independent preparations), in the absence of compounds. ATPase activities are normalized to the respective maximum ATP turnover rate at saturating Ca^2+^ concentration. (B) Concentration-dependent effects of mavacamten, a known cardiac-specific myosin inhibitor. (C) Effect of 50μM compounds on ATPase activities at pCa 4, expressed as percent change in ATPase activity. (D) Effect of 50μM compounds at pCa 8, expressed as percent change in ATPase activity. Error bars indicate standard deviation (n ≥ 8). Significance (*) of difference in effects between skeletal and cardiac was established with Student’s t-test, P<0.05.

To determine the effect of each compound, the ATPase rate of myofibrils in the presence of each compound was normalized to the 1% DMSO only control ATPase rate. The high-throughput NADH-coupled plate reader assay was first validated with a known cardiac-specific myosin-binding compound, mavacamten (8). The effective concentration of mavacamten that gives half-maximal response (EC_50_) is 0.30 ± 0.03 μM for cardiac myofibrils and 7.24 ± 0.96 μM for skeletal (Fig. 5B). Thus, mavacamten selectively inhibits the ATPase activity of cardiac myofibrils compared to skeletal myofibrils, as reported previously (23). Using the same assay, we tested the effect of actin-binding compounds (Table 1) on both types of myofibrils at pCa 4 and pCa 8. Fig. 5C &D summarize the comparative percent change in ATPase activities of respective skeletal and cardiac myofibrils in the presence of 50 μM actin-binding compounds (same concentration as used in actin polymerization). At this concentration, dantrolene was unstable and caused aggregation of myofibrils, and thus was omitted from the ATPase measurement.

At pCa 4 (high calcium), all of the tested compounds act as inhibitors of ATPase for skeletal myofibrils and some compounds had approximately 20% inhibitory effects on cardiac myofibrils (Fig. 5C). Four compounds: honokiol, phenothiazine, tegaserod, and mefloquine show significantly different effects on ATPase of cardiac and skeletal muscle myofibrils under activating conditions. The aforementioned drugs have minimal effect on cardiac myofibril ATPase activity, but they inhibit skeletal myofibrils. Tegaserod exhibits the most significantly different effects on the two muscle systems; it slightly activates cardiac myofibrils and strongly inhibits (30±7 %) skeletal myofibrils. The percent change difference between the two muscle systems for honokiol (13±8 %), phenothiazine (17±8%), and mefloquine (14±8%) were also statistically significant (P<0.05).

Isoform-specific differences in myofibril ATPase activity were also observed at pCa 8 (relaxation). Four compounds, honokiol, fluphenazine, phenothiazine and tegaserod, activated cardiac myofibril ATPase. Of those four, only two also activated skeletal myofibrils: honokiol and tegaserod. Only honokiol, fluphenazine, and phenothiazine showed significantly different effects on the two muscle systems (Fig. 5D). The percent activation for honokiol is 54±13% for cardiac and 20±7% for skeletal. Fluphenazine’s differential effect was less pronounced where cardiac myofibril ATPase is activated by 18±2% while it does not have an effect on skeletal myofibril. Another significant difference in effect on myofibrils ATPase activity at pCa 8 is from Phenothiazine. This drug had minimal inhibitory effect on skeletal myofibril in relaxing conditions, but it increased basal ATPase activity of cardiac myofibril by 33±8 %. Thus, these drugs may play an important role in the calcium sensitization or desensitization of a particular muscle type.

## Discussions

We report a comparative study of a new group of actin-binding compounds, previously detected by our high-throughput and high-precision FRET assay (11), on the polymerization of skeletal and cardiac actin as well as on the ATPase activities of skeletal and cardiac myofibrils, with the goal of testing specificity for actin and muscle isoforms. Our results show that some of these compounds induce isoform-specific effects on actin structure, some of which are reflected in the functional activity of the corresponding muscle type. We conclude that amino acid substitutions between skeletal and cardiac actin induce structural changes in actin that affect each actin’s susceptibility to the effects of bound compounds. Each compound’s mechanism of action probably depends on the structures of the compound and its binding site on actin, resulting in specific changes in the functional interaction of actin and myosin.

### Specificity of compounds for actin isoforms

Actin polymerization is a useful tool to monitor effects of structural changes in actin induced by amino acid substitutions, mutations, or binding to various chemical compounds. Amino acid substitutions in highly conserved actin isoforms result in different polymerization properties of muscle and non-muscle actins (13,24). Similarly, some disease-causing mutations in actin result in altered polymerization kinetics and filament stability (25). Numerous naturally obtained compounds have been found to affect the properties of actin filaments (26–28). Most of these compounds fit into one of two groups: (a) those that destabilize actin filaments (usually by decreasing the rate of polymerization), and (b) those that stabilize actin filaments (usually by increasing the rate of polymerization). In muscle, actin concentration is so high that a significant decrease is the fraction of polymerized (filamentous) actin is not likely. In the present study, we used actin polymerization kinetics to indicate rapidly a significant change in actin’s physical properties. Actin polymerization assays (Fig. 2 and Fig. 3) showed that skeletal and cardiac muscle actin have very similar rates of polymerization as reported earlier (15,29), indicating that the four amino acid substitutions (Glu2Asp, Asp3Glu, Met299Leu and Thr358Ser) between these two isoforms do not have a significant effect on the formation of intermonomer bonds. However, polymerization rates were significantly affected by several of these compounds. Most pronounced changes were observed in the presence of fluphenazine, novantrone, and thioriadazine. These compounds also have pronounced effects on skeletal actin filament dynamics and on the environment of the C-terminus of actin, as detected by transient phosphorescence anisotropy (TPA) in our previous study (11). Additionally, thioriadazine, phenothiazine, and tegaserod exhibit significantly different isoform-specificity. Thus, these compounds are significant modulators of actin structure and dynamics, and their effects show distinct isoform-specificity.

Even though a number of natural small-molecule inhibitors of actin cytoskeleton dynamics have long been recognized as valuable molecular probes for understanding the complex mechanisms of cellular function, few studies have been performed to compare their isoform-specific effects on skeletal and cardiac actin. Phalloidin, a heptapeptide toxin from the poisonous mushroom *Amanita phalloides*, binds tightly and specifically to polymerized actin (30), stabilizing the filaments. A comparison of phalloidin binding to three different isoforms of actin showed that a single amino acid change can alter actin’s affinity for phalloidin (31). Similarly, an actin isoform-specific behavior was also observed for *Clostridium botulinum C2* toxin (32). Thus, a subtle change in actin-compound interaction can change functional behavior. This kind of selectivity must reside in structural differences between the two isoforms. In the present case, the differences in how compounds bind and affect skeletal and cardiac actin must be determined by structural effects of the 4 amino acid substitutions.

### Specificity of compounds for muscle types

Out of all sarcomeric components, small-molecule effectors designed to target and modulate striated and smooth muscle myosin isoforms and troponin C are showing promise in preclinical and clinical trials (7–9) for the treatment of muscle diseases. The balance between contraction and relaxation of muscle is controlled by a shift in calcium concentration. As the mechanical system of the sarcomere is interrelated, any attempt to stabilize the contracted or relaxed state by targeting any one component along the troponin–tropomyosin–actin–myosin continuum, favors the contracted or relaxed state in all components. Thus, an ideal modulator for sarcomeric proteins should work by a delicate shift in calcium sensitivity and should have a minimal effect on downstream regulation. In addition, the compound should be muscle-specific, as desired for its safe clinical application. In our work, we focus on how modulation of actin with actin-binding compounds alters Ca^2+^ dependent muscle regulation as reflected in functional activity. Most of these pharmacologically active compounds affected myofibril ATPase activity of skeletal and cardiac muscle (Fig. 5C &D). Pronounced effects were observed at pCa 4 and at pCa 8 with compounds honokiol, phenothiazine, tegaserod, and mefloquine, showing significant difference on ATPase activity for the two muscle isoforms. Honokiol, phenothiazine and tegaserod act as inhibitors for skeletal myofibril ATPase and as activators for cardiac myofibril ATPase at pCa 4. This isoform-specific behavior probably arises from drug-induced changes in actin structure, as phenothiazine and tegaserod also show isoform-specific behavior on actin polymerization. Fluphenazine, thioridazine, and novantrone inhibited ATPase of both muscle types and had reduced the rate of polymerization of both actin isoforms. This is consistent with our previous study, where these three compounds had maximum effect on F-actin structure and functional interaction with myosin (11). Thus, compound-specific effects are transmitted from actin to the myofibril level. The most interesting features of these compounds were observed at pCa 8, when actin is not actively engaged in interaction with myosin. Several compounds, honokiol, fluphenazine, and phenothiazine showed significant differences in ATPase activity of the two muscle systems at relaxed condition. This suggests that these compounds affect the actin-tropomyosin interface, and induce different structural changes in the actin-tropomyosin complex at low Ca^2+^, exposing an otherwise blocked myosin binding site on actin. Differential behavior of these compounds at pCa 4 and at pCa 8 for a specific muscle also suggests their effect on Ca^2+^ sensitivity for the corresponding muscle.

Therefore, compound-induced structural changes at the actin level affect either the actin-myosin interaction or Ca^2+^ mediated regulation of the corresponding muscle types. A key bottleneck for progressing to clinical trials is the development of compounds that bind with high affinity with the desired target. Our compounds affect actin at μM concentrations, indicating moderate actin binding affinity, which can be optimized in the future through medicinal chemistry. These compounds are already in use as medications for various diseases (Table 1) and some have significant side effects on muscle function (33–39). An effective drug must achieve a balance between specific therapeutic benefit and undesired side effects. Future screening of larger libraries, with the high-throughput structure-based FRET assays and secondary assays developed in this study, will help identify potential actin-binding compounds that can specifically work on a particular muscle type.

## Conclusion

The current study shows that actin-binding compounds, previously discovered by high-throughput FRET screening, have specific biochemical and functional effects that are distinct in cardiac and skeletal actin, and in cardiac and skeletal muscle. This sets the stage to apply the original high-throughput FRET screen to larger small-molecule libraries, leading to the discovery of novel actin-binding compounds with specific therapeutic potential for treating disorders of cardiac or skeletal muscle (11,40).

## Experimental procedures

### Preparation of actin

New Zealand White rabbits (purchased from *Birchwood, Maine USA*) were housed in University Research Animal Resources facility until muscle bundles collection. Rabbits were euthanized by trained veterinarians consistent with the recommendations of the American Veterinary Medical Association (AVMA) Guidelines. Bovine left ventricle was isolated from fresh bovine hearts (*Pel-Freez Biologicals*). Actin was prepared from either rabbit skeletal or bovine cardiac acetone, using the procedure of Pardee and Spudich (41), with slight modifications, as follows. Actin was extracted from acetone powder with cold water and polymerized with 2 mM MgCl_2_, 30 mM KCl and 1 mM ATP for 1.5 hours. Addition of 1 mM ATP at this step ensures full polymerization. Then, 0.6 M KCl was slowly added to polymerized actin and was gently stirred for 1 hr. The polymerized actin was ultracentrifuged at 350, 000xg for 30 minutes at 4°C and the F-actin pellet was suspended in G-Ca buffer (5 mM Tris, 0.5 mM ATP, 0.2 mM CaCl_2_, pH 7.5), followed by clarification at 300,000 x g for 10 minutes. The resultant G-actin was then extensively dialyzed in G-Ca buffer for ~36 hours with periodic changes of G-buffer. G-actin was finally clarified again at 300,000 x g for 10 minutes and protein concentration was measured at 290 nm assuming the molecular weight of 42300 Da and absorption of 0.1% protein 0.63 (42). The prepared actin was used within two days for polymerization experiment.

### Labeling of actin with Pyrene iodoacetamide

Skeletal muscle actin was prepared as described previously (43) by extracting acetone powder of rabbit skeletal muscle with cold water. Actin was polymerized with 30 mM KCl for 1 hour at room temperature, and centrifuged at 350,000xg for 30 minutes at 4°C. The pellet was suspended in G buffer (10 mM Tris, 0.5 mM ATP, 0.2 mM CaCl_2_, pH 7.5). Pyrene-actin was prepared by labeling actin with pyrene iodoacetamide (Invitrogen) as described previously (44), with slight modifications. Actin (24 μM) was polymerized with 0.1 M KCl, 1 mM NaN_3_ and 20 mM Tris (pH 7.5), and the dye, freshly dissolved in DMF, was added at a concentration of 180 μM. After 18 h incubation at 23°C, the labeling was terminated by 10 mM DTT, and actin was ultracentrifuged at 350,000xg for 30 minutes at 4°C. Labeled actin was then resuspended in G-buffer and clarified by 10 min centrifugation at 300,000 x g. Protein concentration was measured by Bradford assay using unlabeled G-actin as a standard.

### Actin polymerization and depolymerization assay

Rabbit skeletal and bovine cardiac actin polymerization rates were determined by the increase in fluorescence caused by incorporation of pyrene-labeled G-actin into actin filaments (45). Pyrene-actin (5 % of total actin concentration) was mixed with globular α-skeletal and α-cardiac actin in G-buffer. Polymerization was induced by the addition of either 2 mM MgCl_2_ or 50 mM KCl to 20 μM G-actin and the increase of pyrene-fluorescence with an excitation wavelength of 350 nm was monitored at 407 nm using a Varian Cary Eclipse spectrofluorometer (total volume of 150 μl) in a microcuvette. Polymerization of actin in the presence of compound was done with a concentration of 50 μM compound with the addition of 2 mM MgCl_2_. For determination of the apparent half-times of polymerization, the kinetic traces were approximated by a single exponential function and the half-times were calculated from the rate constant. Relative change in polymerization was calculated from the ratio of apparent half-times of actin in the presence and in the absence of compound (Fig. 3). For depolymerization, an amount of 20 μM actin containing 5% pyrene actin was first polymerized with 2 mM MgCl_2_ for at least 2 hours at 23°C. F-actin was then diluted 1:20 into G-buffer and the decrease in fluorescence intensity was monitored with the same wavelengths settings as in polymerization (Fig. 4).

### Preparation of skeletal and cardiac myofibrils

Rabbit psoas muscle bundles were isolated and chemically skinned in 0.5% Triton X-100 and stored at −20°C in 50% glycerol with 60 mM KPr, 25 mM MOPS, 2 mM MgCl_2_, 1 mM EGTA, 1 mM NaN_3_, and EDTA-free protease inhibitor cocktail, pH 7.0. Muscles were used within 6 months of the initial skinning process. Skeletal myofibrils were prepared in homogenization solution containing 60 mM KPr, 25 mM MOPS, 1 mM EGTA, and 1 mM NaN_3_, pH 7.0 (18). Muscle bundles were finely cut and subsequently homogenized with a PowerGen 125 Polytron (Fisher Scientific) for 10 minutes. Serial washing via centrifugation at 1,500 x g for 5 min and resuspension with homogenization solution was done to remove residue Triton, glycerol, and protease inhibitors. All steps in the preparation were done at 4°C. Total myofibrillar protein concentration was measured by Bradford assay using BSA as a standard. Freshly prepared myofibrils were used in experiments within 72 hours.

Fresh bovine left ventricle was cut into 1-inch pieces. These chunks were flash-frozen in liquid nitrogen and stored at −80°C until use. Muscle pieces were thawed in base rigor buffer K60 (60 mL KCl, 2 mM MgCl_2_, 20 mM MOPS pH 7.4) containing protease (Sigma#8340) and phosphatase inhibitors (SigmaP0044). Thawed pieces were finely chopped with razor blade and 1 mM EGTA was added for early permeabilization and calcium removal. Muscle bundles were fragmented to myocyte sized fragments with a PowerGen 125 Polytron (Fisher Scientific). Fragments were dounce homogenized prior to skinning in 1.0% Triton X-100 on ice (46). The resulting myofibrils were washed multiple times as described above with K60 to remove Triton, protease and phosphate inhibitors. Finally, myofibril solution was filtered through 70 μm nylon cell strainer and protein concentration was measured by Bradford assay with BSA as a standard. The freshly prepared myofibrils were stored at 4°C that retained near full ATPase activity for about 24 hours.

### Preparation of 96-well assay plates for myofibrillar ATPase assay

The compounds of interest (11) were purchased from TargetMol (Boston, MA USA) at 10 mM stocks in DMSO. The compounds were then distributed into a 96-well template plate and diluted with DMSO to the appropriate concentration for the dose response assays. Final compound concentration was varied from 0-100μM across 12-wells. Final DMSO concentration in each well was kept constant at 1%. Final Assay plates were prepared by transferring 2 μL of compound from the template plates using as Mosquito liquid dispenser from TTP Labtech (Melbourne, UK). These assay plates were stored at −20°C prior to usage.

### Steady State Myofibrillar ATPase Kinetics

An enzyme-coupled, NADH-linked ATPase assay was used to measure skeletal and cardiac myofibril ATPase activity in 96-well microplates at 23°C. Ionic strength was maintained below 0.1 M, so myofilaments remain insoluble and stable (47). Skeletal myofibril was contained in 50 mM MOPS, 35 mM KCl, 2 mM MgCl_2_, 1 mM EGTA, pH 7.0. Cardiac myofibril was preserved in 20 mM MOPS, 35 mM NaCl, 5 mM MgCl_2_, 1 mM EGTA, pH 7.0. Ionic strength dependence difference between two the myofibrils led us to using slightly different buffer condition for each muscle system. Each well in microplate contained an assay mix of 0.84 mM phosphoenolpyruvate, 0.17 mM NADH, 10 U/vol pyruvate kinase, 20 U/vol lactate dehydrogenase. The concentration of free Ca^2+^ was controlled by EGTA buffering (48). Myofibrils were dispensed in microplate and were incubated with assay mix plus compound of interest for 20 minutes at 23°C. The concentration of myofibrils used were 0.01 mg/ml for skeletal and 0.05 mg/ml for cardiac. The assay was started upon the addition of ATP, at a final concentration of 2.1 mM (total volume to 200 μL), and absorbance was recorded at 340 nm in a SpectraMax Plus microplate spectrophotometer from Molecular Devices (Sunnyvale, CA). The steady-state myofibrillar ATPase rate was measured from NADH oxidation, measured as the rate of decrease in absorption at 340 nm for 30 minutes, with absorption data collected every 15 seconds. All data and statistical analysis of steady-state kinetics were conducted with the OriginPro program. The results were fitted with the Hill equation:

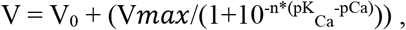

where V is the ATPase rate and n is the Hill coefficient.

All ATPase data are normalized to a per second per myosin head scale, assuming that 50% of the myofibril content is myosin (49). Average data are presented as mean ± standard deviation. Sample means are derived from eight or more experimental repeats.

The effects of drugs on ATPase activity of myofibrils were determined in the presence of 50 μM compound, as prepared above, at activating pCa of 4.0 and relaxing pCa of 8.0 (Fig. 5). The effect of 500 nM thapsigargin (a SERCA modulator) on myofibrils ATPase activity inhibition was tested across a 12-points pCa range. In all dose response experiments, a 1% DMSO only control was included.

## Data availability

All data discussed are presented within the article.

## Acknowledgements

Fluorescence experiments and drug plate preparations were performed at the Biophysical Technology Center, University of Minnesota. We thank Chandini Nair and Ananya Tripathi for technical assistance with earlier experiments.

## Funding and additional information

This work was supported by NIH grants to DDT (R01AR032961, R37AG26160) and LAP (T32GM008244).

## Conflict of Interest

DDT holds equity in, and serves as President of, Photonic Pharma LLC. This relationship has been reviewed and managed by the University of Minnesota. Photonic Pharma had no role in this study. The authors declare no conflicts of interest with the contents of this article.

## Author Contributions

PG and DDT designed the research. PG, LAP, SL and AW prepared samples, performed experiments, and analyzed the data. PG, LAP, EP and DDT wrote the paper.

